# Baseline microbiome composition impacts resilience to and recovery following antibiotics

**DOI:** 10.1101/2024.03.31.587491

**Authors:** Chia-Yu Chen, Ulrike Löber, Hendrik Bartolomaeus, Lisa Maier, Dominik N. Müller, Nicola Wilck, Víctor Hugo Jarquín-Díaz, Sofia K. Forslund-Startceva

## Abstract

The gut microbiome of healthy individuals naturally undergoes temporal changes linked to the dynamics of its community components^1^. These dynamics are only observable in longitudinal studies; they are particularly relevant to understanding ecosystem responses to external environment disturbances. External exposures, such as antibiotic treatment, significantly reshape the gut microbiome, impacting both pathogen and commensal microbes^2^. The gut microbiome plays pivotal roles in digestion, nutrient absorption, and mental health, influencing immune systems, obesity, and various diseases^3-6^. Consequently, beyond the short-term effects on the host gut microbiome dynamics, alterations resulting from antibiotic exposure also have enduring repercussions on human health and physiological equilibrium^7^. Therefore, enhancing gut microbiome resilience during antibiotic treatment is essential, with the goal of mitigating prolonged adverse effects. Here, we explored the impact of pre-antibiotic microbial and functional profiles on resilience, suggesting that specific baseline features exhibit greater resilience to antibiotics-induced changes. Our results identified an uncultured *Faecalibacterium prausnitzii* taxon as a species at baseline associated with diminished resilience. We demonstrated that this association could be linked to the role of this *F. prausnitzii* taxon as a keystone species. Additionally, we observed the influence of other commensal bacteria, such as *Bifidobacterium animalis* and *Lactobacillus acidophilus*, as well as functional modules, such as multidrug resistance efflux pump, on resilience. This lays the foundations for designing targeted strategies to promote a resilient gut microbiome before antibiotic treatment, alleviating possible prolonged effects on human health.

---

Antibiotics are known to disrupt the balance of the gut microbiome and induce long-term microbiome alterations, often termed dysbiosis in the literature. Altered microbiome compositions have been linked to various health conditions like inflammatory bowel disease, obesity, and cardiovascular disease, especially the loss of producers of beneficial metabolites like short-chain fatty acids (SCFAs)^8,9^. SCFAs play a beneficial role in health maintenance and disease development by interacting with the host immune system^10^. Post-antibiotic dysbiosis is characterized by diminished microbial diversity, the absence of crucial taxa, and metabolic shifts^11^. Even brief antibiotic usage, especially in early childhood, can induce lasting changes in the gut microbiome^12^. Probiotic consumption alongside or following antibiotic treatment has been proposed to alleviate antibiotic-induced disruptions. However, the existing literature lacks a consensus, presenting conflicting results^13-17^. Recognizing the profound effects of antibiotics on the gut microbiome is crucial to understanding the factors influencing resilience to mitigate long-term consequences and maintain overall host health.

Resilience encompasses two aspects: resistance, the ability to withstand changes upon external exposure, and recovery, the rate or time to revert to the original state (Figure 1A)^18^. Factors influencing resilience include host immune status, microbiome diversity, and functional redundancy^19^. Functional redundancy illustrates that diverse organisms across different taxa perform the same functional role in the ecosystem^20^. High functional redundancy indicates that altering species diversity may have a limited effect on overall ecosystem functionality^21^. Here, we hypothesize that the microbial or functional composition before antibiotic treatment plays a pivotal role in resilience.

**Figure 1.**
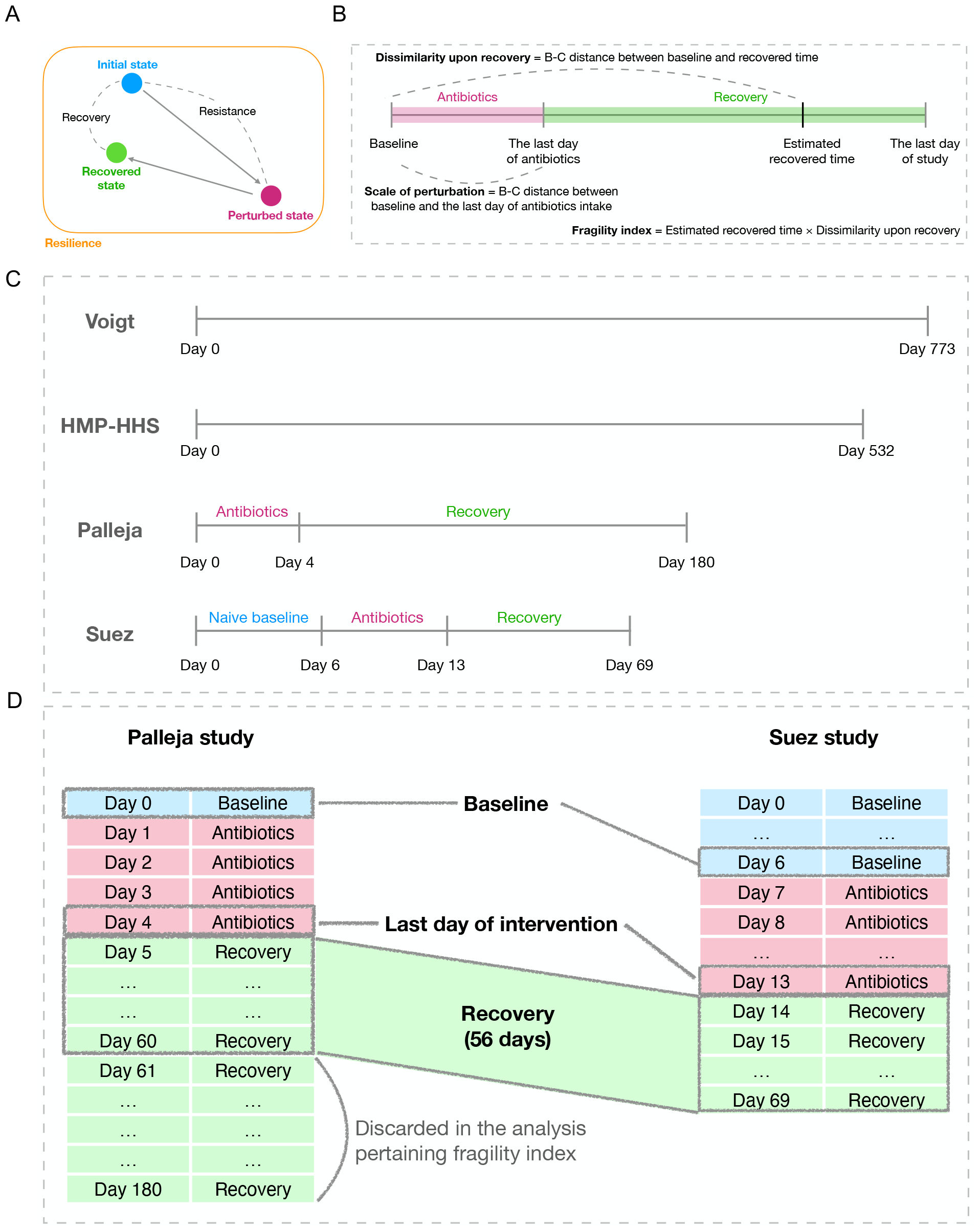
Study concept and design. **(A)** The illustration shows the concept of resistance and resilience upon antibiotics treatment. Resilience is represented as how much a microbial community change compared with its original state after the antibiotics perturbation, while recovery can be measured by how much the post-perturbation state resembles the initial state. **(B)** The figure shows the components of the fragility index, which includes the scale of perturbation, estimated recovered time and dissimilarity upon recovery. B-C is the abbreviation of Bray-Curtis. **(C)** The data included in this study are illustrated. The sample size of each study is as follows. Voigt (n = 7, including the subject “Alien”); HMP-HHS (n = 55); Palleja (n = 12); Suez before antibiotics treatment (n = 21); Suez after antibiotics treatment (n = 16). The discrepancy between the sample size of Suez before/after antibiotics is because some of the individuals underwent further post-antibiotics intervention (i.e., probiotics supplement or autologous fecal transplant) and were excluded here. **(D)** The illustration shows the alignment of time points between the Palleja and the Suez studies. Day 0 in the Palleja and day 6 in the Suez studies are taken as the baseline. Day 4 in the Palleja and day 13 in the Suez studies are the last day of intervention. The Palleja study is curtailed to day 60 so that its recovery duration is the same as that of Suez (56 days).

To test whether the effect of antibiotics and the recovery rate were associated, we introduced the term “fragility index” (FI). The FI reflects how vulnerable a microbial community is during exposure to external disruptions (Figure 1B). The FI is computed as follows:

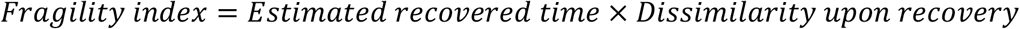

We assessed the *estimated recovered time* to the time in which the Bray-Curtis distance is the closest to zero against the baseline^22^. The *dissimilarity upon recovery* corresponds to the Bray-Curtis distance between the baseline and the estimated recovered time. The FI depends on the resistance and the recovery aspects; a higher fragility index indicates a less resilient microbial community when exposed to external perturbations. Henceforth, we refer to “resistance” as the “scale of perturbation”, defined as the Bray-Curtis distance between before and after the antibiotic treatment. In this study, we first examined the contrast in microbial community dynamics between unperturbed states and perturbation caused by antibiotic intervention. Subsequently, we tested the hypothesis that microbial or functional profiles are crucial in determining resilience.

To quantify the microbial perturbation induced by antibiotics compared with unperturbed microbial community dynamics in healthy human individuals, we collected, reanalyzed, and compared published datasets. Our study incorporated a total of four datasets. Two datasets, from Voigt et al.^23^ (hereafter Voigt) and the Human Microbiome Project Healthy Human Subjects^24^ (hereafter HMP-HHS), correspond to healthy human cohorts tracked for temporal variation in their gut microbiome. The other two datasets, from Palleja et al.^25^ (hereafter Palleja) and Suez et al.^13^ (hereafter Suez), correspond to antibiotic intervention studies (Figure 1C). In the Voigt study, only one participant was under antibiotic treatment. The subject, referred to as “Alien”, received ceftriaxone for four days due to infection and was segregated from other participants for the analysis. The Palleja and Suez studies involved healthy human cohorts treated with antibiotics for four and seven days, respectively. Both studies used a cocktail of broad-spectrum antibiotics and vancomycin (Extended Table 1). The antibiotic combinations ensure comparability and suitability for a combined analysis. The Palleja and Suez studies lack continuous (day-by-day) taxonomic and functional abundance data. Therefore, we employed natural cubic spline interpolation to estimate taxonomic and functional abundances at the missing time points and then calculate the estimated recovered time^26^. Moreover, the Palleja and Suez studies have different intervention and follow-up time points. While the Suez study has a longer antibiotic intervention duration, the Palleja study has more extended follow-up time points. Hence, their respective time points were realigned to ensure comparability in the estimated recovery time between both studies (Figure 1D).

To assess the impact of antibiotics on the microbial community in comparison with the natural dynamics of a healthy microbiota, we computed the Bray-Curtis distance between each time point and the baseline for all individuals (Figure 2A, Extended Figure 1A). We employed polynomial regression to model the Bray-Curtis dissimilarity over time for antibiotic-treated and non-treated individuals. In the non-treated group, a shift occurred, reaching stability and forming a plateau around 0.4 over time (Figure 2B, Extended Figure 1B). For the antibiotic-treated group, the Bray-Curtis distance increased to 0.8 following antibiotic exposure and gradually returned to a level similar to that of the non-treated group. Notably, from day 300 to 400, the antibiotic-treated group’s distance from the baseline showed no significant difference compared with the non-treated group, as indicated by overlapping confidence intervals. This shows that antibiotic exposure induced a perturbation in the gut microbiota, reaching about twice the scale of the shift in healthy individuals, and eventually returning to a level similar to the non-treated group during recovery. We observed a significant correlation between the fragility index and the scale of perturbation based on both taxonomic (genus, Spearman’s correlation p < 0.05, rho = 0.49) and functional (KEGG module, Spearman’s correlation p < 0.05, rho = 0.49) profiles (Figure 2C, 2D). This indicates that the scale of perturbation serves as a suitable proxy for the fragility index, and, in turn, is associated with overall resilience. Moreover, the scales of perturbation between taxa and functional profiles are correlated (Extended Figures 1C, 1D).

**Figure 2.**
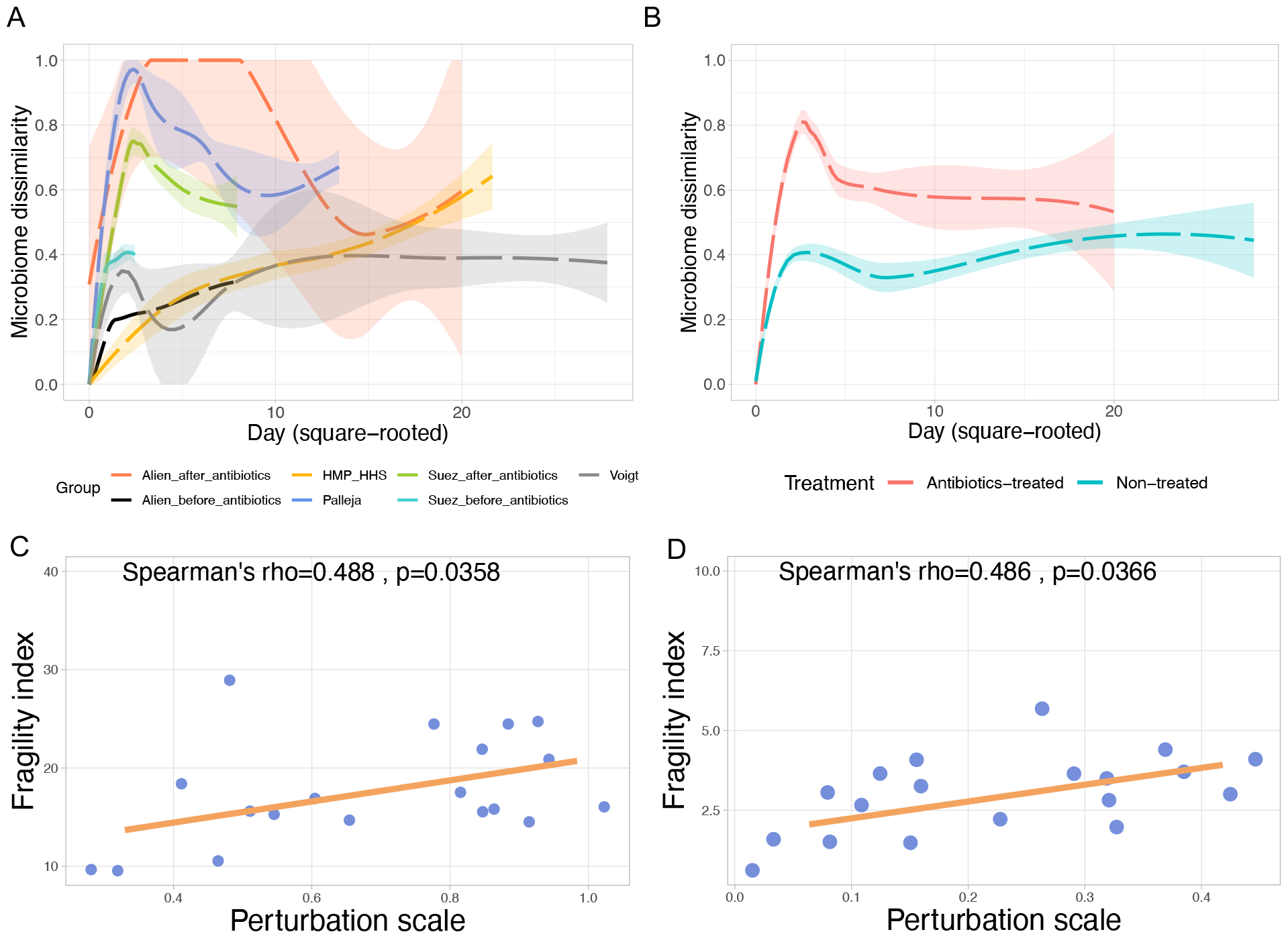
Gut microbiome shift is larger in antibiotics-treated group, and the scale of perturbation is correlated with the fragility index. **(A, B)** The line plots show the microbiome dissimilarity (Bray-Curtis distance) at species level between the baseline and each time point at subsequent days. Samples are grouped by (A) different data source, which are Alien_before_antibiotics (n = 1), Alien_after_antibiotics (n = 1), HMP-HHS (n = 55), Palleja (n = 12), Suez_before_antibiotics (n = 21), Suez_after_antibiotics (n = 16) and Voigt (n = 6), or (B) different treatment, which are non-treated (n = 83) and antibiotics-treated (n = 29). Smooth curves (dashed lines) were fitted to show the overall pattern in each group. Transparent shades indicate 95% confidence interval. The discrepancy between the sample size of Suez before/after antibiotics is because some of the individuals underwent further post-antibiotics intervention (i.e., probiotics supplement or autologous fecal transplant) and were excluded here. In (B), the significance of treatment was shown by a linear model fitting the microbiome dissimilarity as a function of treatment*day(square rooted). P-values for treatment and the interaction term treatment:day were both < 0.001 **(C, D)** Scatter plots showing the correlation between the perturbation scale and the fragility index are based on (C) microbial genus and (D) KEGG modules. Spearman’s correlation test results are annotated on the plots. The orange solid lines are the linear regression fitted to the points to show the overall trend.

Next, our aim was to characterize the microbiome and functional profiles that are resilient upon antibiotic perturbation. First, we discovered a negative correlation (Spearman’s p < 0.01, rho = -0.64) between functional redundancy and the scale of perturbation (Figure 3A). Then, we applied the Random Forest (RF) method, an ensemble learning technique that combines independently sampled decision trees to improve model robustness and generalization performance^27^. Utilizing RF regression models, we tested which microbial and functional features were linked to antibiotic perturbation and detected an uncultured *F. prausnitzii* as the most relevant predictor (Spearman’s rho = 0.6, Figure 3B). Our RF results were validated by correlation to leave-one-out cross-validation (LOOCV) (Extended Figure 2). In contrast, baseline Shannon diversity and the baseline abundance of *Blautia* sp., *Ruminococcus* sp. CAG 254, *Prevotella* sp. CAG 520, *L. acidophilus*, and *B. animalis* are negatively correlated with the scale of perturbation (Spearman’s rhos < -0.53, Figure 3B). At the genus level (Figure 3B column 2), baseline Shannon diversity (Spearman’s rho = -0.56) and the abundance of Dehalococcoidales gen. incertae sedis (Spearman’s rho = -0.56) are significantly and negatively correlated with the scale of perturbation. We built a second RF model to predict the scale of perturbation based on the baseline functional profile, including KEGG modules and gut metabolic modules (GMMs) (Figure 3B columns 3, 4)^28,29^. We found that multidrug resistance efflux pumps (KEGG M00645 and M00700) are among the most influential ones, contributing to functional stability. Given the relevance of a known butyrate producer like *F. prausnitzii* to predict the scale of perturbation^30^, it is interesting to detect that the baseline abundance of modules associated with the production of this SCFA like 4-aminobutyrate degradation (GMM MF0042) and crotonyl-coA from succinate (GMM MF0115) negatively correlated with the scale of perturbation^31^. We observed that *Bifidobacterium, Lactobacillus*, and MF0042 are consistent features across different models (Figure 3B, all columns). Specific functions, for example, multidrug resistance efflux pumps M00646 and M00647 in Figure 3B column 5 show positive correlations with the scale of perturbation, while M00645 and M00700 in Figure 3B column 3 exhibit a negative correlation. It is important to recognize that, although classified as multidrug resistance efflux pumps, these systems can exhibit inherent differences in their composition and functional mechanisms^32^. When we test which taxonomic or functional features are the best predictors of the FI, our model identified fewer significant correlations, as shown in Extended Figures 3 and 4 (e.g., only six features showed correlation significance in Extended Figure 3).

**Figure 3.**
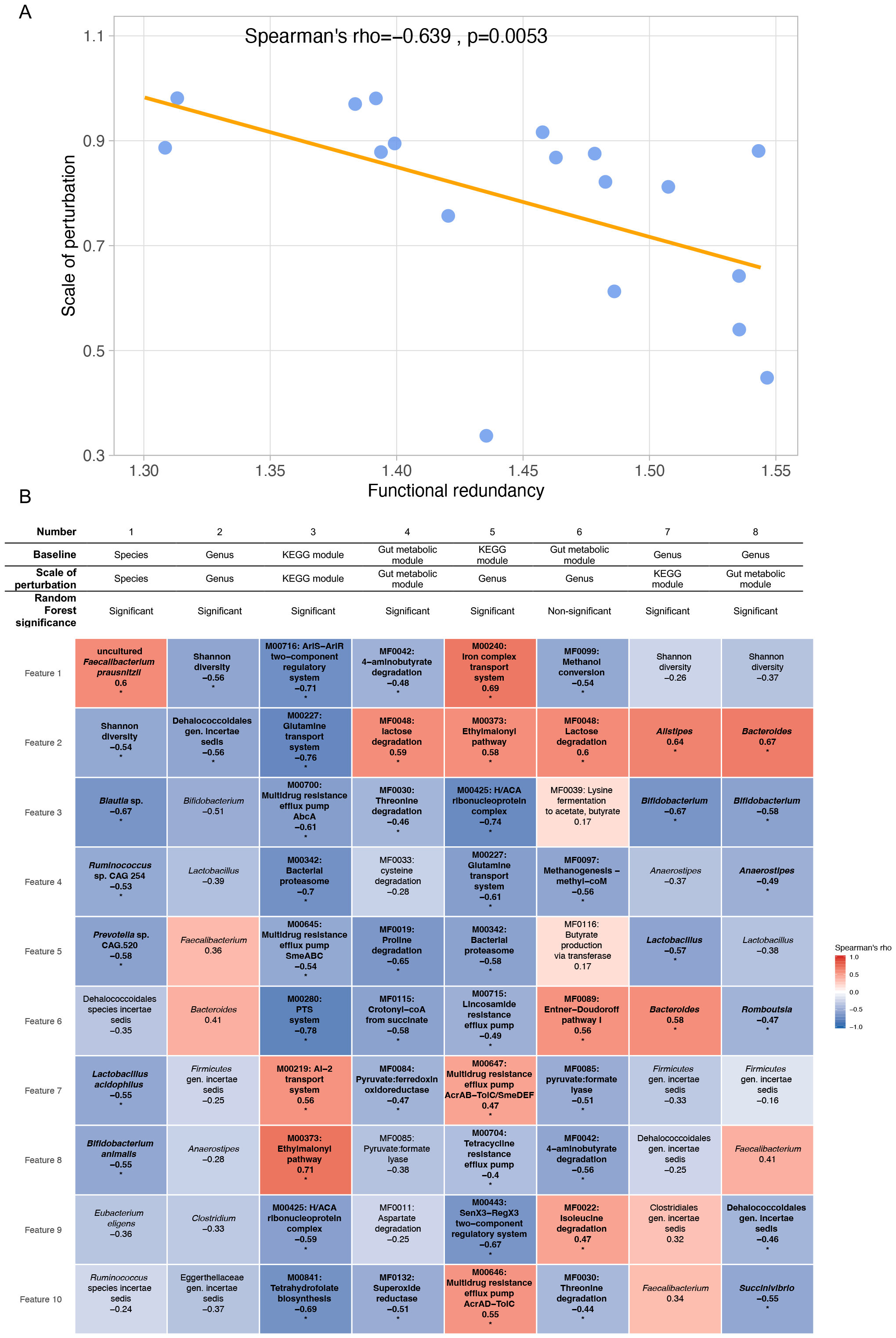
Characterization of the resilient microbiome and functional profile. **(A)** The scatter plot shows the correlation between baseline functional redundancy and the microbiome dissimilarity (Bray-Curtis distance) of the gut microbiome at genus level between the baseline and post-antibiotics treatment. The functional redundancy is calculated as the Shannon diversity of the species-specific variants (GMGC) within each gene family (KEGG KO). Spearman’s correlation test’s p-value and rho of are shown in the figure. The orange solid line is the linear regression fitted to the points to show the overall trend. **(B)** This heatmap shows the result of RF regression fitting the scale of perturbation. The random forest models were trained using 5-fold cross-validation. Each column represents the scale of perturbation based on one feature predicted by the other baseline feature. RF significance is based on the permutation (n = 1000) result, where the negative MAE of each RF is compared to the permutated negative MAE. If a negative MAE is larger than > 95 % of the permutated values, then it is regarded as significant (p < 0.05). Colors represent Spearman’s correlation between the feature with the scale of perturbation. Text below the feature name is the Spearman’s rho value. Asterisks and bold texts denote that Spearman’s correlation q < 0.1. Benjamini-Hochberg procedure was used to adjust the p values within each column (n = 10).

**Figure 4.**
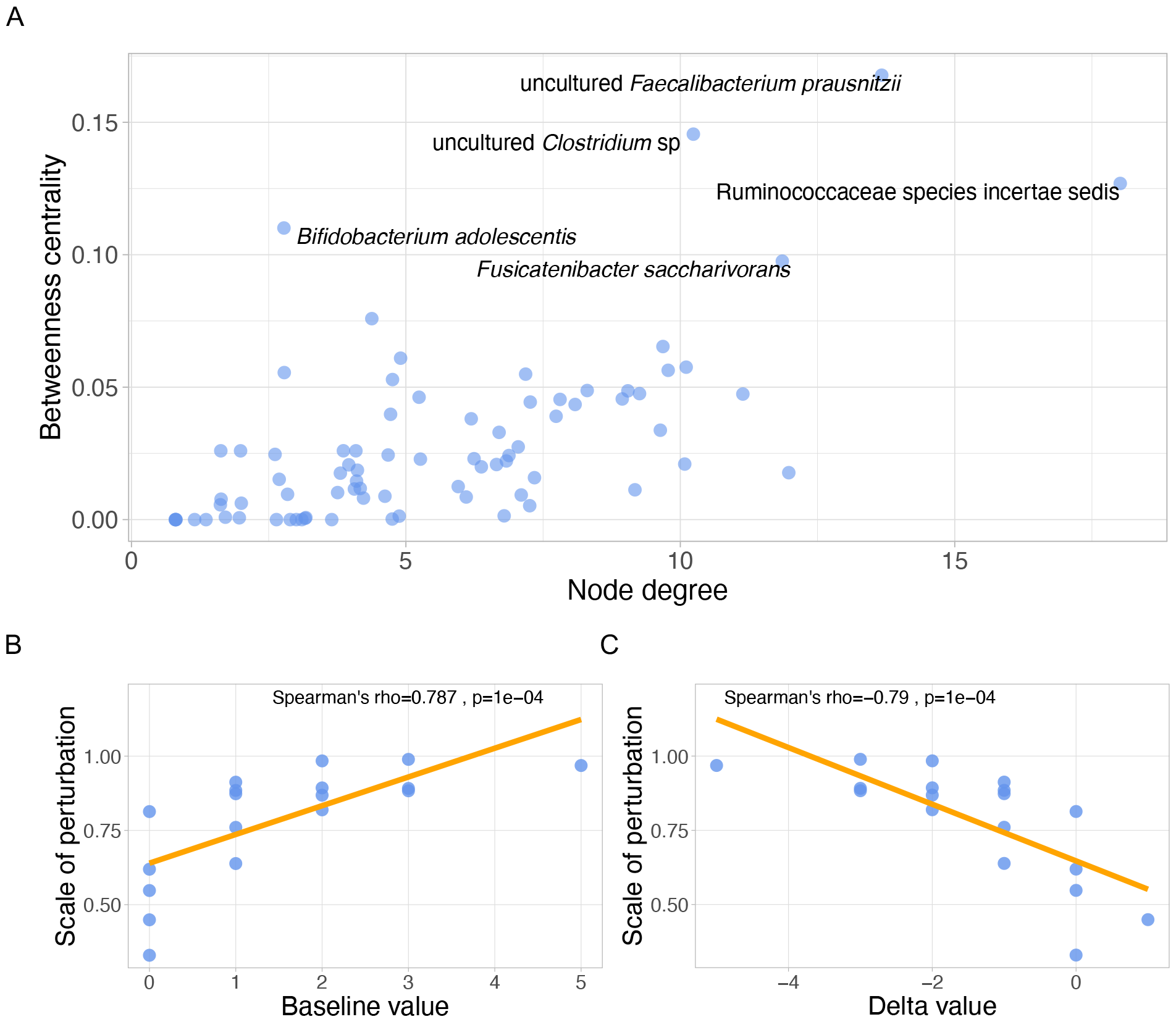
Uncultured *F. prausnitzii* taxon as a keystone species. **(A)** The scatter plot shows the betweenness centrality against node degree of each species, which is the result from co-occurrence network analysis. **(B, C)** These figures plot the scale of perturbation against the (B) the abundance of *F. prausnitzii* 06108 at the baseline and (C) the difference in *F. prausnitzii* 06108 abundance between before and after antibiotics treatment (after – before). Spearman’s correlation test’s p-value and rho of are shown in the figures. The orange solid lines are the linear regression fitted to the points to show the overall trend.

In addition to the uncultured *F. prausnitzii* taxon, other recognized SCFA producers like *B. animalis, L. acidophilus, Blautia* spp., and *Ruminococcus* sp. CAG 254 were identified as relevant taxonomic features to predict the effect of antibiotic perturbation^33,34^. We concluded that the majority of SCFA-producing species at baseline play a protective role in preserving microbial communities against extensive perturbation induced by antibiotic treatment. Based on the relevance of uncultured *F. prausnitzii* in our models, we speculated about its potential role as a keystone species within the microbial community. A keystone species exerts a disproportionately significant influence on the other community members relative to its abundance^35^. To test whether the uncultured *F. prausnitzii* has a significant role in the microbial community, we constructed a co-occurrence network based on significant taxa-taxa correlations (Spearman’s correlations, Benjamini-Hochberg corrected q < 0.1) (Extended Figure 5) and used centrality measures to characterize the relevance of this taxon. In network analyses, nodes with elevated betweenness centrality (i.e., the frequency of a node acting as a bridge along the shortest path between two other nodes) may signify key connectors, while nodes with high node degrees (i.e., the number of connections a node has to other nodes) may represent hubs^36^. Both high betweenness centrality and node degree serve as indicators for potential keystone species. We observed that uncultured *F. prausnitzii* possessed the highest network betweenness centrality (Figure 4A), confirmed by an alternative method (Extended Figures 6A, 6B). We observed a depletion of the uncultured *F. prausnitzii* after the antibiotic treatment (Spearman’s correlation p < 0.01, rho = -0.78) and a decrease in the abundances of all its associated taxa (n=14) based on the network analysis (Extended Figure 5) (Spearman’s correlations, Benjamini-Hochberg corrected q < 0.05, rho values between -0.40 ∼ -0.69). We suggest that the decline in uncultured *F. prausnitzii* during antibiotic treatment is linked to the reduction of other associated species, consistent with a role as a keystone species.

The uncultured *F. prausnitzii* taxon corresponds to two metagenomic Operational Taxonomic Units (mOTUs), ref-mOTU-v25-06108 (hereafter referred to as *F. prausnitzii* 06108) and ref-mOTU-v25-06110 (hereafter referred to as *F. prausnitzii* 06110). To discern the relevance of these mOTUs, we ran new RF regression models using baseline mOTUs (restricted to those associated with significant species in Figure 3 column 1) to predict the scale of perturbation. Notably, our results highlight *F. prausnitzii* 06108 as the foremost influential feature, positioning it as the most crucial predictor, while *F. prausnitzii* 06110 holds a lower rank (8^th^) in importance. We observed that the baseline abundance of *F. prausnitzii* 06108 positively correlated with the scale of perturbation, while its delta value (before/after antibiotics) negatively correlated with the scale of perturbation (Figure 4B, 4C). The latter suggests that an increased loss of *F. prausnitzii* 06108 post-antibiotics leads to a larger scale of perturbation. Our findings propose that the depletion of *F. prausnitzii* 06108 during antibiotic treatment precipitates the loss of other associated species. Furthermore, we examined the variations in functional profiles within the two mOTUs, delineating their distinctions using KEGG modules, pathways, and BRITE, and identified several disparate functions (Extended Figures 7–9). For example, *F. prausnitzii* 06108 was exclusively correlated with KEGG modules such as cytochrome c oxidase, cytochrome bd ubiquinol oxidase and acylglycerol degradation, and KEGG pathways such as protein digestion and absorption.

Our study explores the impact of antibiotics on the gut microbiome. Our results indicate that antibiotic-induced perturbation in the studied datasets is approximately twice the scale of natural drift in healthy individuals over the same time interval. Even in the absence of any external source of perturbation such as antibiotics, drift over time, as well as measurement variability, results in an average within-donor gut microbiome similarity over time of around 0.4 Bray-Curtis distance. This is in agreement with previous findings in a study involving fecal samples from healthy Belgian women collected over 6 weeks, where within-individual Bray-Curtis distances fell between 0.25 and 0.51. We detected a negative correlation between functional redundancy and the scale of perturbation, consistent with previous studies suggesting that high redundancy contributes to stable functioning in a microbiome community under external perturbation^37^. We found a positive correlation between functional redundancy and the extent of perturbation upon antibiotics exposure and established the fragility index as a proxy to comprehensively describe resilience in a microbiome context. When we explored the predictive potential of baseline microbiome abundance, we detected several commensal SCFA producers associated with the scale of perturbation. In particular, an *F. prausnitzii* taxon was anti-correlated with resilience, consistent with a keystone species role of this commensal, which we confirmed by co-occurrence network analysis, as previously reported^38^. Conversely, *Blautia* sp., *Ruminococcus* sp., *L. acidophilus*, and *B. animalis* at baseline were associated to higher resilience. The same applies to baseline Shannon diversity, in line with previous literature^19^. Lastly, several functional modules, including multidrug resistance efflux pumps and butyrate-producing pathways, emerged as predictors of the degree of community perturbation upon antibiotic exposure.

Our findings complement the conventional notion of employing probiotics based on *Bifidobacterium* or *Lactobacillus* during or after antibiotics treatment to facilitate recovery, suggesting other potential candidates for supplementation post-antibiotics. Even within the same species, we found that different strains may impact community resilience differently. Future efforts will focus on validating the resilience-enhancing capabilities of SCFA producers through experimental studies. Our results suggest a potential role of probiotics before antibiotic treatment to fortify gut microbiome resilience, which future experiments should seek to validate.

## Methods

### Data acquisition

Raw metagenomic sequencing data from the Palleja^25^, Suez^13^, and Voigt^23^ studies were acquired from the European Nucleotide Archive (ENA) database, using the accession numbers ERP022986, PRJEB28097, and ERP009422, respectively. Metadata for these studies was obtained from the corresponding publications. For the HMP-HHS study^24^, sequencing data were obtained from the Human Microbiome Project (HMP) portal at https://portal.hmpdacc.org/projects/t. The metadata for this study were controlled-access and were retrieved from the database of Genotypes and Phenotypes (dbGaP). The individuals in the HMP_HHS study who had taken antibiotics were excluded from our analysis.

### Shotgun metagenomic processing

The metagenomic shotgun sequences were subjected to processing using NGLess v1.3^39^. Reads underwent quality filtering, with a minimum read length of 45 bp and a minimum Phred quality score of 25. Sequences meeting these criteria were subsequently mapped to the human genome (adapted from hg38), with a minimum requirement of a 45 bp match and at least 90% identity, and those mapping to the human genome were filtered out. Taxonomic classification was determined by aligning the non-human reads to metagenomic Operational Taxonomic Units (mOTUs v2.5)^40^ using bwa with default parameters. Functional profiling was done by aligning non-human reads to the Global Microbial Gene Catalog (GMGC v1.0)^41^. Instances where reads mapped to multiple genes in the reference database were addressed utilizing the “all1” method. Next, the GMGC IDs were binned into KEGG KOs^28^, and then further into KEGG modules or gut metabolic modules (GMM)^29^. To mitigate the impact of differences in sequencing depth in downstream analysis, rarefaction on abundance tables was performed using RTK tool v0.93.1^42^.

### Abundance interpolation

To obtain continuous data (i.e., data for every single day) from the discrete raw data, natural cubic spline interpolation was employed for both taxonomic and functional abundance data^26,43^. This technique was utilized to estimate abundances at missing time points by constructing piece wise third-order (cubic) polynomials that pass through the known data points. The spline interpolation was performed using the R function splinefun(method = “natural”).

### Timepoint realignment

To ensure uniformity in the estimated recovery time between the Suez and Palleja studies, their respective time points underwent realignment. Day 0 in Palleja and day 6 in Suez were designated as the baseline. Day 4 in Palleja and day 13 in Suez were regarded as the final days of intervention. Ultimately, follow-up time points were extended up to day 60 in Palleja to generate estimates of recovery time that align with the time frame of Suez.

### Alpha and beta diversities analysis

Both Shannon diversity and Bray-Curtis distance were calculated using the vegan v2.5 R package^44^. The rarefied abundance table at both the species and genus levels was utilized for Bray-Curtis distance computation.

### Functional redundancy

To assess functional redundancy, characterized by the presence of multiple species-specific variants within gene families, the following method was employed. A Shannon metric was computed for each individual based on the GMGC profiles corresponding to each gene family (e.g., KEGG KO). This calculation was conducted independently for each gene family. Subsequently, the average Shannon metric was determined by aggregating the values across all gene families for each individual.

### Correlation analysis

Spearman’s correlation tests were done using the R basic function cor.test(method = “spearman”). These included the correlation between taxonomic or functional profile abundances and scale of perturbation or fragility index, as well as all other instances in the figures that explicitly specified Spearman’s correlation test. Benjamini-Hochberg correction of the p values was done using the function p.adjust(method = “fdr”) in base R.

### Random forest regression

Random forest regression was carried out using the Python machine-learning package scikit-learn^45^. First, the out-of-bag error (OOB error) was employed to select optimal hyperparameters, including the maximum number of features, maximum depth, and number of estimators. Subsequently, two separate evaluations were conducted: one using 5-fold cross-validation and the other using leave-one-out cross-validation (LOOCV). The evaluation metric for both procedures was negative mean absolute error (negative MAE)^46^. Following this, permutation (n = 1000) was performed by randomly shifting the dependent variable against the independent variables, and all resulting negative MAEs were recorded. The distribution of these permutated negative MAEs was analyzed to assess the overall performance of the random forest regression.

### Keystone species analysis

Spearman’s correlations were computed for all species using R. Subsequently, only the statistically significant correlations (Benjamini-Hochberg corrected q < 0.1) were employed in constructing a co-occurrence network. The betweenness centrality and node degree were then calculated. Cytoscape v3.9.1^47^ was employed for network construction, visualization, and the computation of betweenness centrality and node degree. Additionally, for independent validation, the cooccur R package^48^, which applies the probabilistic model to assess species co-occurrence, was also employed.

### Linking functional profiles to specific mOTUs for the uncultured *F. prausnitzii* taxon

From the mOTU v2.5 database, the uncultured *F. prausnitzii* fraction comprises two mOTUs, namely ref_mOTU_v25_06108 and ref_mOTU_v25_06110. Genomic data and GMGC profiling for the uncultured *F. prausnitzii* were extracted, and GMGC IDs with low abundance (sum across samples < 7) were filtered out. Spearman correlation analyses were then conducted between the abundances of the two target *F. prausnitzii* mOTUs and each GMGC ID, followed by Benjamini-Hochberg (BH) correction. Correlations between GMGC and mOTUs were filtered based on significance (q < 0.05). GMGC IDs were binned into KEGG modules, pathways, or BRITEs categories using the KEGG database (https://www.kegg.jp). NA values, indicating no significant correlation of GMGC belonging to a KEGG module, pathway, or BRITE with one mOTU (but significant correlation with the other), were replaced with 0. The focus was on positive correlations between GMGC and mOTUs, excluding negative correlations, to identify co-abundant mOTUs and genes within the context of the observed uncultured *F. prausnitzii* taxon.

### Visualization

Figures were generated using ggplot2 v3.3.5^49^ and the ggpubr v0.4.0 (https://cran.r-project.org/package=ggpubr) R packages (https://rdocumentation.org/packages/ggpubr/). Smooth curves in Figures 2A and 2B are local polynomial regressions fitted by using the function ggplot2::geom_smooth(method = “loess”). In Figures 2C, 2D, 3A, 4B, and 4C, as well as Extended Figures 1C and 1D, the smooth lines were derived from linear regression and fitted using ggplot2::geom_smooth(method = “lm”).

## Data availability

For raw sequencing files, please see “data acquisition” in Methods section. Processed data are available at https://github.com/CCY-dev/Microbiome_resilience_data_code.

## Code availability

Scripts are available at https://github.com/CCY-dev/Microbiome_resilience_data_code.

## References

1 Vandeputte, D. et al. Temporal variability in quantitative human gut microbiome profiles and implications for clinical research. Nature Communications 12, 6740 (2021). 10.1038/s41467-021-27098-7

2 Macfarlane, S. Antibiotic treatments and microbes in the gut. Environ Microbiol 16, 919–924 (2014). 10.1111/1462-2920.12399

3 Ogunrinola, G. A., Oyewale, J. O., Oshamika, O. O. & Olasehinde, G. I. The Human Microbiome and Its Impacts on Health. International Journal of Microbiology 2020, 8045646 (2020). 10.1155/2020/8045646

4 Clapp, M. et al. Gut microbiota’s effect on mental health: The gut-brain axis. Clin Pract 7, 987 (2017). 10.4081/cp.2017.987

5 Meyer, K. et al. Association of the Gut Microbiota With Cognitive Function in Midlife. JAMA Network Open 5, e2143941–e2143941 (2022). 10.1001/jamanetworkopen.2021.43941

6 Sun, J. & Chang, E. B. Exploring gut microbes in human health and disease: Pushing the envelope. Genes Dis 1, 132–139 (2014). 10.1016/j.gendis.2014.08.001

7 Francino, M. P. Antibiotics and the Human Gut Microbiome: Dysbioses and Accumulation of Resistances. Front Microbiol 6, 1543 (2015). 10.3389/fmicb.2015.01543

8 Carding, S., Verbeke, K., Vipond, D. T., Corfe, B. M. & Owen, L. J. Dysbiosis of the gut microbiota in disease. Microbial Ecology in Health and Disease 26, 26191 (2015). 10.3402/mehd.v26.26191

9 Morris, G. et al. The Role of the Microbial Metabolites Including Tryptophan Catabolites and Short Chain Fatty Acids in the Pathophysiology of Immune-Inflammatory and Neuroimmune Disease. Molecular Neurobiology 54, 4432–4451 (2017). 10.1007/s12035-016-0004-2

10 Xiong, R.-G. et al. Health Benefits and Side Effects of Short-Chain Fatty Acids. Foods 11, 2863 (2022).

11 Duan, H. et al. Antibiotic-induced gut dysbiosis and barrier disruption and the potential protective strategies. Critical Reviews in Food Science and Nutrition 62, 1427–1452 (2022). 10.1080/10408398.2020.1843396

12 Lange, K., Buerger, M., Stallmach, A. & Bruns, T. Effects of Antibiotics on Gut Microbiota. Digestive Diseases 34, 260–268 (2016). 10.1159/000443360

13 Suez, J. et al. Post-Antibiotic Gut Mucosal Microbiome Reconstitution Is Impaired by Probiotics and Improved by Autologous FMT. Cell 174, 1406–1423 e1416 (2018). 10.1016/j.cell.2018.08.047

14 Engelbrektson, A. et al. Probiotics to minimize the disruption of faecal microbiota in healthy subjects undergoing antibiotic therapy. Journal of Medical Microbiology 58, 663–670 (2009). 10.1099/jmm.0.47615-0

15 Dahiya, D. & Nigam, P. S. Antibiotic-Therapy-Induced Gut Dysbiosis Affecting Gut Microbiota&mdash;Brain Axis and Cognition: Restoration by Intake of Probiotics and Synbiotics. International Journal of Molecular Sciences 24, 3074 (2023).

16 Videlock, E. J. & Cremonini, F. Meta-analysis: probiotics in antibiotic-associated diarrhoea. Alimentary Pharmacology & Therapeutics 35, 1355–1369 (2012). 10.1111/j.1365-2036.2012.05104.x

17 Chng, K. R. et al. Metagenome-wide association analysis identifies microbial determinants of post-antibiotic ecological recovery in the gut. Nature Ecology & Evolution 4, 1256–1267 (2020). 10.1038/s41559-020-1236-0

18 Medeiros, L. P., Song, C. & Saavedra, S. Merging dynamical and structural indicators to measure resilience in multispecies systems. J Anim Ecol 90, 2027–2040 (2021). 10.1111/1365-2656.13421

19 Dogra, S. K., Doré, J. & Damak, S. Gut Microbiota Resilience: Definition, Link to Health and Strategies for Intervention. Front Microbiol 11, 572921 (2020). 10.3389/fmicb.2020.572921

20 Eisenhauer, N., Hines, J., Maestre, F. T. & Rillig, M. C. Reconsidering functional redundancy in biodiversity research. npj Biodiversity 2, 9 (2023). 10.1038/s44185-023-00015-5

21 Li, L. et al. Revealing proteome-level functional redundancy in the human gut microbiome using ultra-deep metaproteomics. Nature Communications 14, 3428 (2023). 10.1038/s41467-023-39149-2

22 Kers, J. G. & Saccenti, E. The Power of Microbiome Studies: Some Considerations on Which Alpha and Beta Metrics to Use and How to Report Results. Front Microbiol 12, 796025 (2021). 10.3389/fmicb.2021.796025

23 Voigt, A. Y. et al. Temporal and technical variability of human gut metagenomes. Genome Biol 16, 73 (2015). 10.1186/s13059-015-0639-8

24 Human Microbiome Project, C. Structure, function and diversity of the healthy human microbiome. Nature 486, 207–214 (2012). 10.1038/nature11234

25 Palleja, A. et al. Recovery of gut microbiota of healthy adults following antibiotic exposure. Nat Microbiol 3, 1255–1265 (2018). 10.1038/s41564-018-0257-9

26 RH, B., J.C., B. & B.A., B. Hermite and Cubic Spline Interpolation. An Introduction to Splines for Use in Computer Graphics and Geometric Modelling. (Morgan Kaufmann, 1998).

27 Breiman, L. Random Forests. Machine Learning 45, 5–32 (2001). 10.1023/A:1010933404324

28 Kanehisa, M., Furumichi, M., Sato, Y., Kawashima, M. & Ishiguro-Watanabe, M. KEGG for taxonomy-based analysis of pathways and genomes. Nucleic Acids Research 51, D587–D592 (2022). 10.1093/nar/gkac963

29 Vieira-Silva, S. et al. Species–function relationships shape ecological properties of the human gut microbiome. Nature Microbiology 1, 16088 (2016). 10.1038/nmicrobiol.2016.88

30 Ferreira-Halder, C. V., Faria, A. V. d. S. & Andrade, S. S. Action and function of Faecalibacterium prausnitzii in health and disease. Best Practice & Research Clinical Gastroenterology 31, 643–648 (2017). 10.1016/j.bpg.2017.09.011

31 Vital, M., Howe, A. C. & Tiedje, J. M. Revealing the bacterial butyrate synthesis pathways by analyzing (meta)genomic data. mBio 5, e00889 (2014). 10.1128/mBio.00889-14

32 Huang, L. et al. Bacterial Multidrug Efflux Pumps at the Frontline of Antimicrobial Resistance: An Overview. Antibiotics 11, 520 (2022).

33 Markowiak-Kopeć, P. & Śliżewska, K. The Effect of Probiotics on the Production of Short-Chain Fatty Acids by Human Intestinal Microbiome. Nutrients 12, 1107 (2020). 10.3390/nu12041107

34 Usta-Gorgun, B. & Yilmaz-Ersan, L. Short-chain fatty acids production by Bifidobacterium species in the presence of salep. Electronic Journal of Biotechnology 47, 29–35 (2020). 10.1016/j.ejbt.2020.06.004

35 Power, M. E. et al. Challenges in the Quest for Keystones: Identifying keystone species is difficult—but essential to understanding how loss of species will affect ecosystems. BioScience 46, 609–620 (1996). 10.2307/1312990

36 Tipton, L. et al. Fungi stabilize connectivity in the lung and skin microbial ecosystems. Microbiome 6, 12 (2018). 10.1186/s40168-017-0393-0

37 Shade, A. Microbiome rescue: directing resilience of environmental microbial communities. Current Opinion in Microbiology 72, 102263 (2023). 10.1016/j.mib.2022.102263

38 Tudela, H., Claus, S. P. & Saleh, M. Next Generation Microbiome Research: Identification of Keystone Species in the Metabolic Regulation of Host-Gut Microbiota Interplay. Frontiers in Cell and Developmental Biology 9 (2021). 10.3389/fcell.2021.719072

39 Coelho, L. P. et al. NG-meta-profiler: fast processing of metagenomes using NGLess, a domain-specific language. Microbiome 7, 84 (2019). 10.1186/s40168-019-0684-8

40 Milanese, A. et al. Microbial abundance, activity and population genomic profiling with mOTUs2. Nature Communications 10, 1014 (2019). 10.1038/s41467-019-08844-4

41 Mende, D. R. et al. proGenomes2: an improved database for accurate and consistent habitat, taxonomic and functional annotations of prokaryotic genomes. Nucleic Acids Research 48, D621–D625 (2019). 10.1093/nar/gkz1002

42 Saary, P., Forslund, K., Bork, P. & Hildebrand, F. RTK: efficient rarefaction analysis of large datasets. Bioinformatics 33, 2594–2595 (2017). 10.1093/bioinformatics/btx206

43 Steinway, S. N., Biggs, M. B., Loughran, T. P., Jr., Papin, J. A. & Albert, R. Inference of Network Dynamics and Metabolic Interactions in the Gut Microbiome. PLoS Comput Biol 11, e1004338 (2015). 10.1371/journal.pcbi.1004338

44 Dixon, P. Vegan, a package of R functions for community ecology. J Veg Sci 14, 927–930 (2003). DOI 10.1111/j.1654-1103.2003.tb02228.x

45 Pedregosa, F. et al. Scikit-learn: Machine Learning in Python. J. Mach. Learn. Res. 12, 2825–2830 (2011).

46 Hodson, T. O. Root-mean-square error (RMSE) or mean absolute error (MAE): when to use them or not. Geosci. Model Dev. 15, 5481–5487 (2022). 10.5194/gmd-15-5481-2022

47 Shannon, P. et al. Cytoscape: a software environment for integrated models of biomolecular interaction networks. Genome research 13, 2498–2504 (2003). 10.1101/gr.1239303

48 Griffith, D. M., Veech, J. A. & Marsh, C. J. cooccur: Probabilistic Species Co-Occurrence Analysis in R. Journal of Statistical Software, Code Snippets 69, 1–17 (2016). 10.18637/jss.v069.c02

49 Wickham, H. ggplot2: Elegant Graphics for Data Analysis., (Springer-Verlag, 2016).

