## Extended figures for "Baseline microbiome composition impacts resilience to and recovery following antibiotics"

**Extended Table 1.** Antibiotics used in the studies.

| Antibiotics | Classification | Study |
| --- | --- | --- |
| Ceftriaxone | Cephalosporin | Alien (from Voigt) |
| Meropenem | Carbapenem | Palleja |
| Vancomycin | Glycopeptide | Palleja |
| Gentamicin | Aminoglycoside | Palleja |
| Ciprofloxacin | Fluoroquinolone | Suez |
| Metronidazole | Nitroimidazole | Suez |

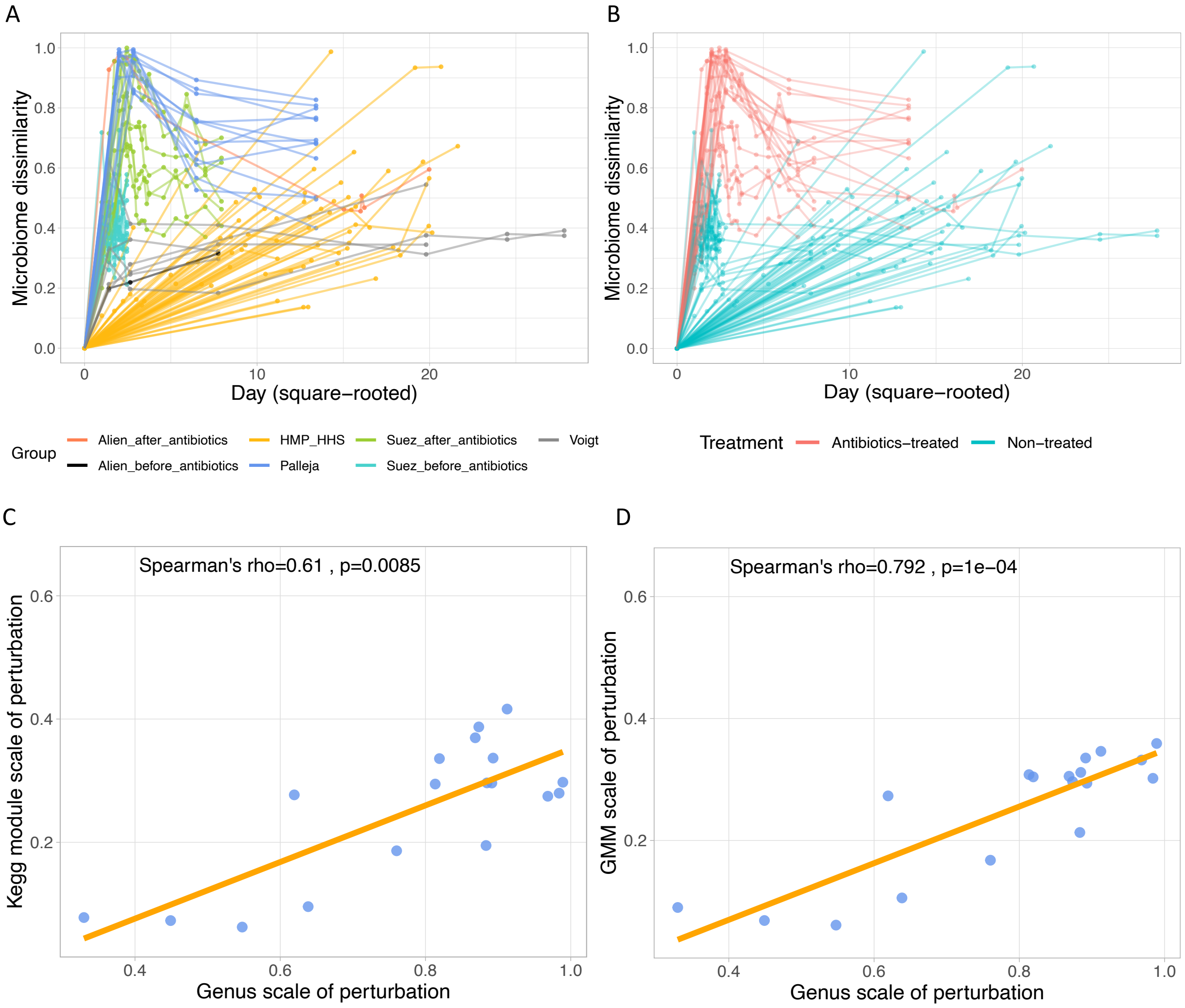


**Extended figure 1. Gut microbiome shift is larger in antibiotics-treated group, and the scale of perturbation between taxa and functional profile are correlated.** The plots show the microbiome dissimilarity (Bray-Curtis distance) at the species level between the baseline and each time point at subsequent days. Samples are colored by **(A)** different data source, which are Alien_before_antibiotics (n = 1), Alien_after_antibiotics (n = 1), HMP-HHS (n = 55), Palleja (n = 12), Suez_before_antibiotics (n = 21), Suez_after_antibiotics (n = 16) and Voigt (n = 6), or **(B)** different treatment, which are non-treated (n = 83) and antibiotics-treated (n = 29). Each point denotes one sample. Lines connect points of the same individual. The discrepancy between the sample size of Suez before/after antibiotics is because some of the individuals underwent further post-antibiotics intervention (i.e., probiotics supplement or autologous fecal transplant) and were excluded here. (C, D) The correlation between the scales of perturbance of genus and (C) KEGG module and (D) gut metabolic module (GMM). The orange solid lines are the linear regression fitted to the points to show the overall trend.

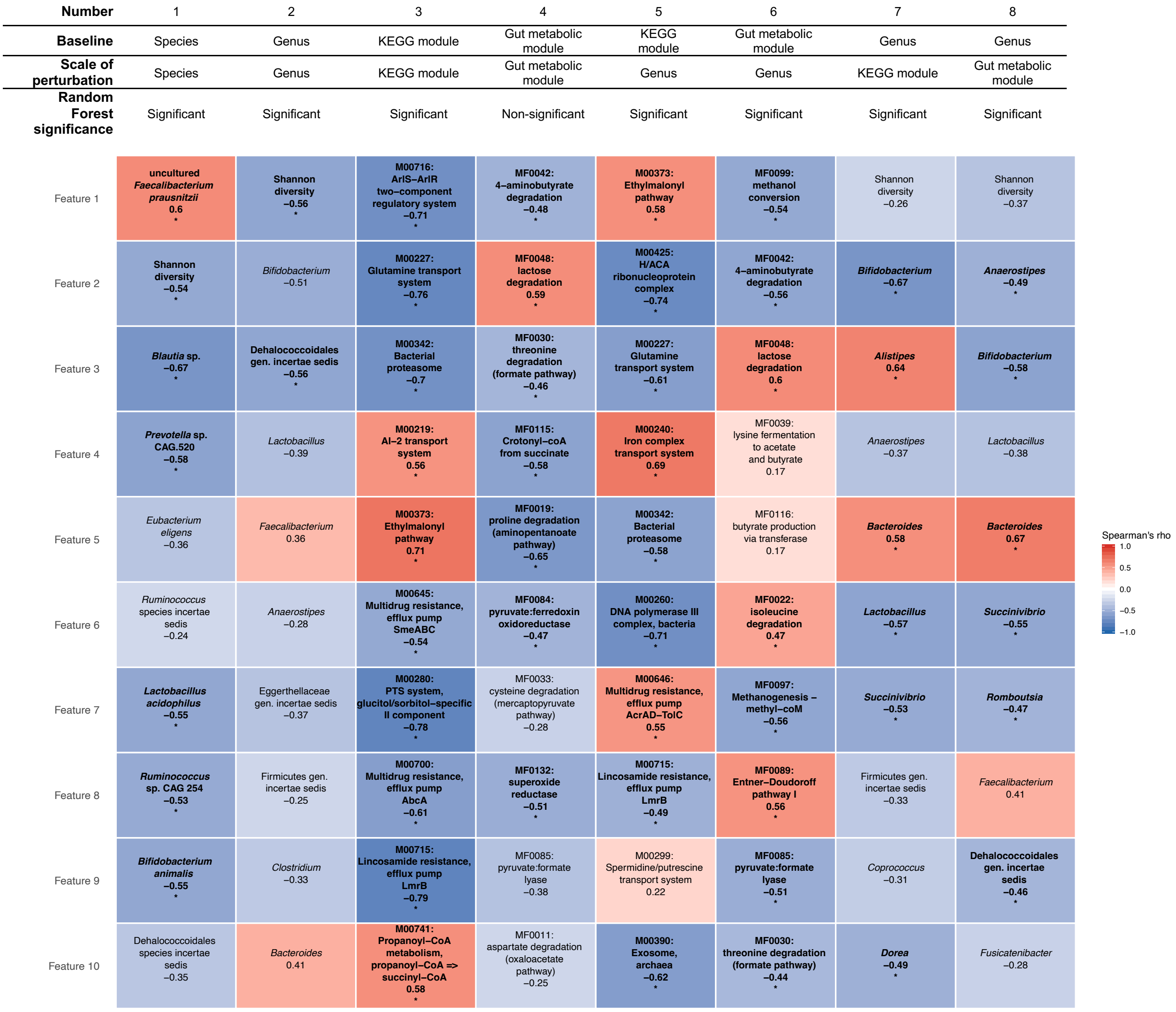


**Extended figure 2. The result of random forest regression fitting the scale of perturbation.** The random forest models were trained using leave-one-out cross-validation (LOOCV). Each column represents the scale of perturbation based on one feature predicted by the other baseline feature. RF significance is based on the permutation (n = 1000) result, where the negative MAE of each RF is compared to the permutated negative MAE. If a negative MAE is larger than > 95 % of the permutated values, then it is regarded as significant ($p < 0.05$). Colors represent Spearman's correlation between the feature with the scale of perturbation. Text below the feature name is the Spearman's rho value. Asterisks and bold texts denote that Spearman's correlation $q < 0.1$. Benjamini-Hochberg procedure was used to adjust the p values within each column (n = 10).

| Number | 1 | 2 | 3 | 4 |
|---|---|---|---|---|
| **Baseline** | Species | Genus | KEGG module | Gut metabolic module |
| **Fragility index** | Species | Genus | KEGG module | Gut metabolic module |
| **Random forest significance** | Non-significant | Non-significant | Non-significant | Non-significant |
| Feature 1 | *Prevotella copri* −0.42 | *Eubacterium* −0.41 | **M00212: Ribose transport system −0.61** * | **MF0019: proline degradation (aminopentanoate pathway) −0.66** * |
| Feature 2 | *Prevotella* sp. incertae sedis −0.18 | Lachnospiraceae gen. incertae sedis 0.41 | **M00841: Tetrahydrofolate biosynthesis −0.52** * | MF0020: valine degradation 0.44 |
| Feature 3 | ***Roseburia intestinalis* 0.64** * | *Prevotella* −0.07 | M00383: ECV complex 0.39 | **Perturbation scale 0.64** * |
| Feature 4 | *Ruminococcus* sp. incertae sedis −0.47 | Perturbation scale 0.49 | M00523: RegB−RegA (redox response) two−component regulatory system 0 | MF0103: nitrate reduction (assimilatory) 0.4 |
| Feature 5 | *Bacteroides caecimuris* 0.39 | Shannon diversity −0.34 | **M00790: Pyrrolnitrin biosynthesis, tryptophan => pyrrolnitrin −0.58** * | MF0091: beta−D−glucuronide and D−glucuronate degradation 0.38 |
| Feature 6 | uncultured *Faecalibacterium prausnitzii* 0.28 | *Coprococcus* −0.4 | M00261: DNA polymerase alpha / primase complex −0.21 | MF0008: tyrosine degradation (hydroxyphenylacet−aldehyde pathway) 0.07 |
| Feature 7 | *Ruminococcus bicirculans* 0.21 | Ruminococcaceae gen. incertae sedis 0.37 | M00806: PTS system, maltose−specific II component −0.38 | MF0076: arabitol degradation −0.37 |
| Feature 8 | Lachnospiraceae sp. incertae sedis 0.51 | *Barnesiella* 0.13 | M00217: D−Allose transport system −0.34 | MF0011: aspartate degradation (oxaloacetate pathway) 0.01 |
| Feature 9 | *Bifidobacterium ruminantium* 0.27 | *Holdemanella* −0.46 | M00333: Type IV secretion system 0.26 | MF0072: ribitol degradation −0.41 |
| Feature 10 | *Faecalibacterium* sp. 0.28 | *Bifidobacterium* −0.32 | M00202: Oligogalacturonide transport system 0.39 | MF0097: Methanogenesis – methyl−coM −0.47 |

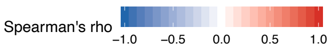


**Extended figure 3. The result of random forest regression fitting the fragility index.** The random forest models were trained using 5-fold cross-validation. Each column represents the fragility index based on one feature predicted by the other baseline feature. RF significance is based on the permutation (n = 1000) result, where the negative MAE of each RF is compared to the permutated negative MAE. If a negative MAE is larger than > 95 % of the permutated values, then it is regarded as significant ($p < 0.05$). Colors represent Spearman's correlation between the feature with the fragility index. Text below the feature name is the Spearman's rho value. Asterisks and bold texts denote that Spearman's correlation $q < 0.1$. Benjamini-Hochberg procedure was used to adjust the p values within each column (n = 10).

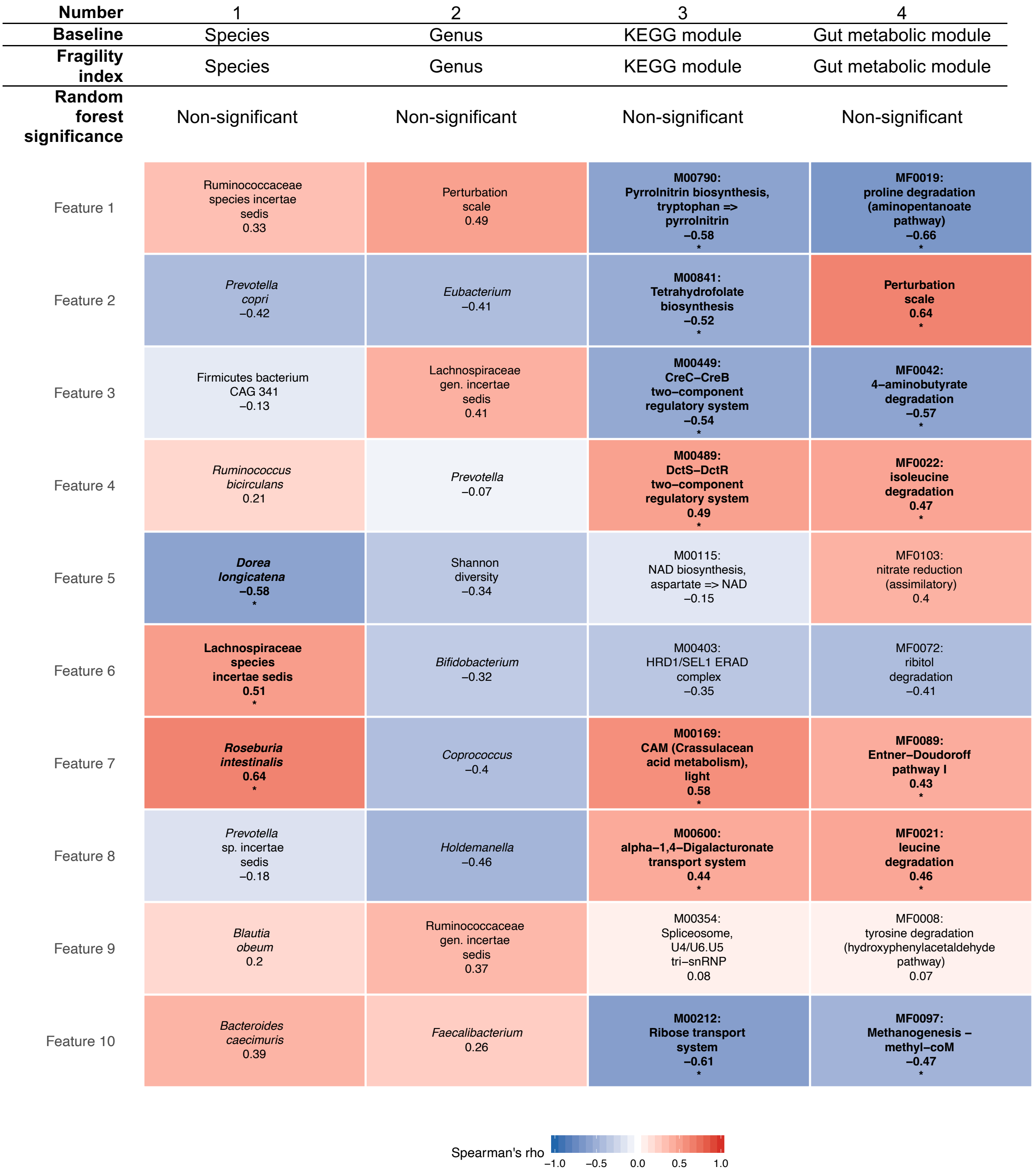


**Extended figure 4. The result of random forest regression fitting the fragility index.** The random forest models were trained using leave-one-out cross-validation (LOOCV). Each column represents the fragility index based on one feature predicted by the other baseline feature. RF significance is based on the permutation (n = 1000) result, where the negative MAE of each RF is compared to the permutated negative MAE. If a negative MAE is larger than > 95 % of the permutated values, then it is regarded as significant ($p < 0.05$). Colors represent Spearman's correlation between the feature with the fragility index. Text below the feature name is the Spearman's rho value. Asterisks and bold texts denote that Spearman's correlation $q < 0.1$. Benjamini-Hochberg procedure was used to adjust the p values within each column (n = 10).

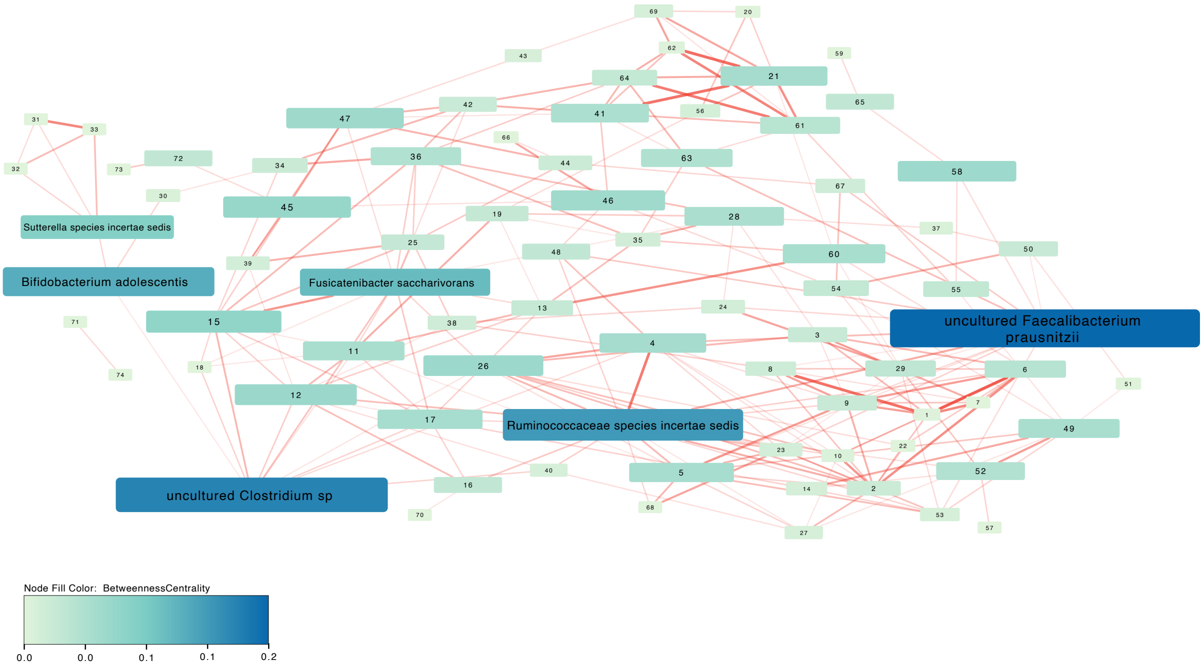


1 *Alistipes shahii*
2 *Bacteroides rodentiumuniformis*
3 *Bacteroides stercoris*
4 Clostridiales species incertae sedis
5 *Faecalibacterium* sp
6 Firmicutes sp
7 *Parabacteroides merdae*
8 *Paraprevotella clara*
9 *Alistipes putredinis*
10 *Roseburia hominis*
11 *Anaerostipes hadrus* uncultured
12 *Blautia obeum/wexlerae*
13 *Coprococcus comes*
14 *Eubacterium* species incertae sedis
15 Lachnospiraceae species incertae sedis
16 *Oscillibacter* sp
17 *Anaerostipes hadrus*
18 *Blautia obeum* uncultured
19 *Clostridium* sp
20 *Ruminococcus* sp CAG 254
21 *Succinivibrio dextrinosolvens*
22 *Bacteroides* sp
23 *Oscillibacter* sp ER4
24 *Roseburia intestinalis*
25 *Ruminococcus torques*
26 *Bacteroides dorei/vulgatus*
27 *Odoribacter splanchnicus*
28 *Blautia massiliensis*
29 *Firmicutes* bacterium CAG 341
30 *Bifidobacterium ruminantium*
31 *Bifidobacterium pseudocatenulatum*
32 *Lactobacillus gasseri/paragasseri*
33 *Bifidobacterium catenulatum/kashiwanohense*
34 *Eubacterium ramulus*
35 *Dorea formicigenerans*
36 *Dorea longicatena*
37 *Roseburia inulinivorans*
38 *Bacteroides massiliensis*
39 Clostridiales sp
40 *Eubacterium eligens* Candidatus
41 *Dialister* sp CAG 486
42 *Prevotella* sp CAG 520
43 *Romboutsia timonensis*
44 *Roseburia faecis*
45 *Eubacterium hallii*
46 *Eubacterium* sp CAG 180
47 *Blautia* sp
48 *Eubacterium eligens*
49 *Clostridium* sp CAG 138
50 *Roseburia* species incertae sedis
51 *Clostridium* sp CAG 127
52 *Clostridium* species incertae sedis
53 *Faecalibacterium prausnitzii*
54 *Ruminococcus bromii*
55 *Ruminococcus* sp CAG 177
56 Bacteroidales species incertae sedis
57 *Roseburia* sp
58 *Eggerthella* species incertae sedis
59 *Eubacterium rectale*
60 *Ruminococcus* species incertae sedis
61 *Holdemanella biformis*
62 *Mitsuokella jalaludinii*
63 *Prevotella copri*
64 *Prevotella* species incertae sedis
65 *Collinsella aerofaciens*
66 *Butyricicoccus* species incertae sedis
67 *Faecalibacterium* species incertae sedis
68 *Dialister invisus*
69 Firmicutes species incertae sedis
70 *Parabacteroides distasonis*
71 *Firmicutes* sp/ *Eubacterium* sp CAG 76
72 *Streptococcus thermophilus*
73 *Streptococcus parasanguinis*
74 *Ruminococcus* sp/ *Eubacterium* sp CAG 76 incertae sedis

**Extended figure 5. Co-occurrence network at species level.** Spearman's correlations between all species were calculated, and then those significant ones (Benjamini-Hochberg corrected q < 0.1) were used to construct a co-occurrence network. Red and blue represent positive and negative correlation, respectively.

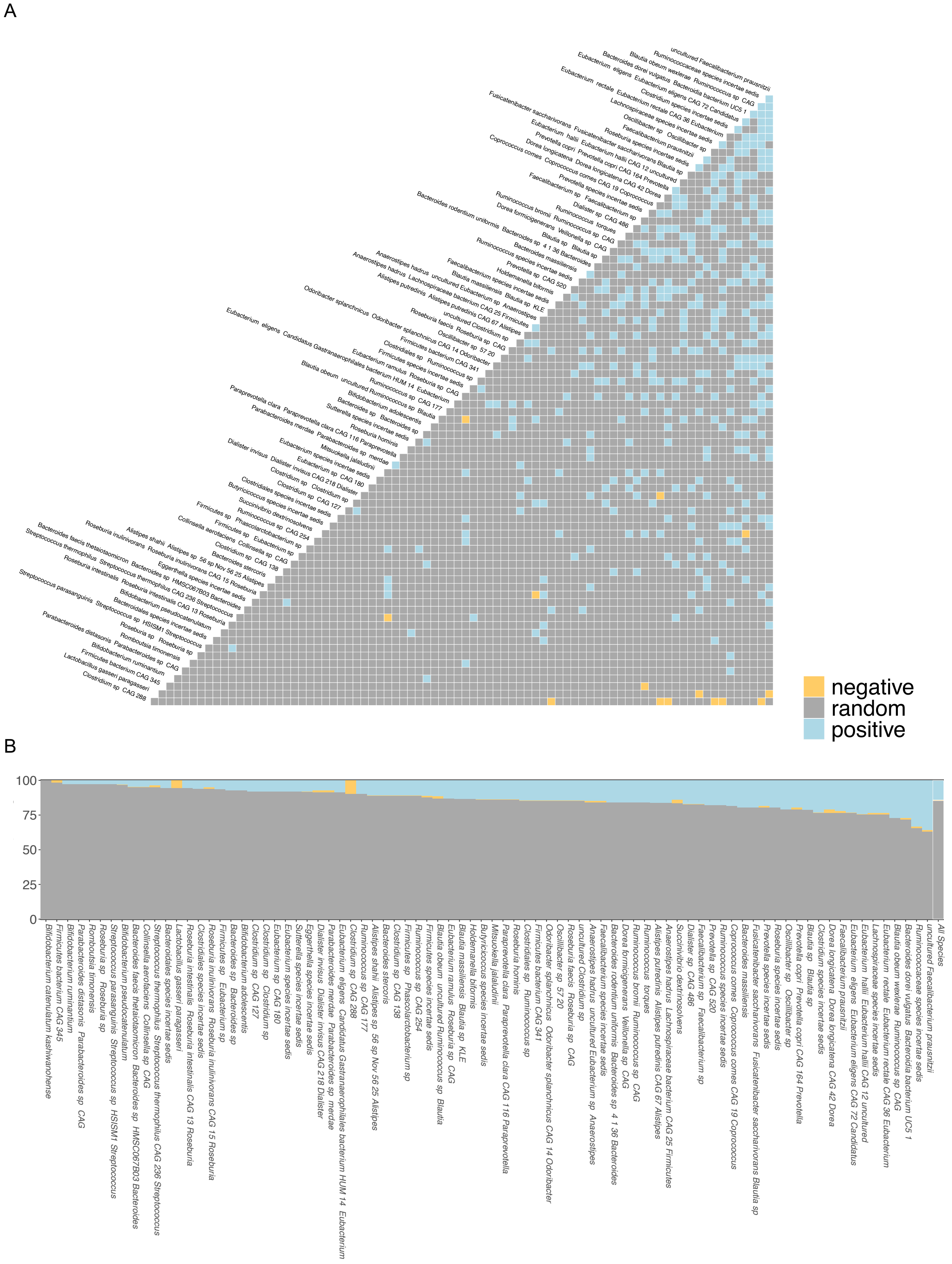


**Extended figure 6. Species co-occurrence results of the microbiome community (A)** Species co−occurrence Matrix **(B)** Species association profile. This analysis was done using the cooccur R package.

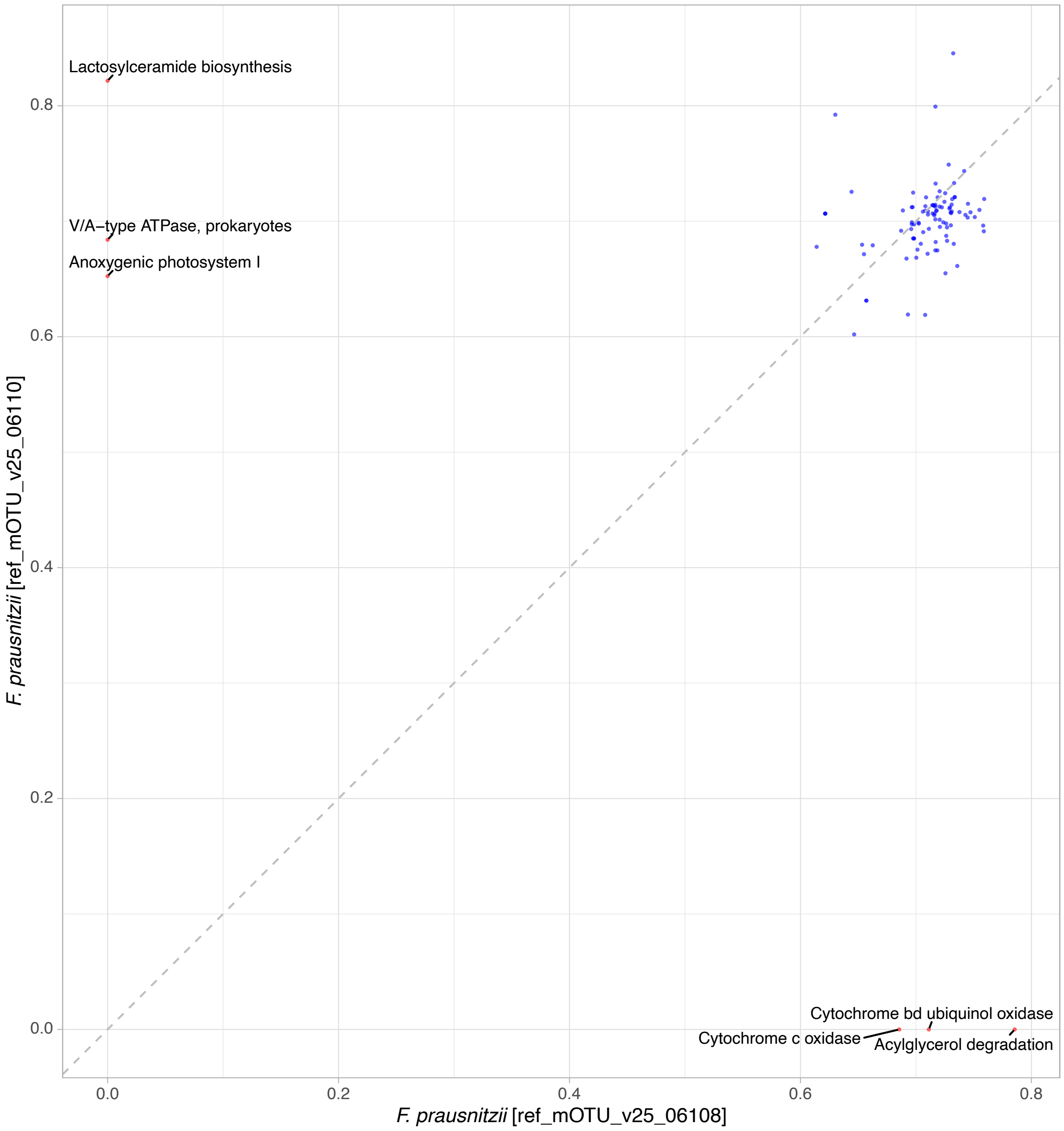


**Extended figure 7. The KEGG modules that are correlated with the two uncultured *F. prausnitzii* mOTUs.** The x and y axes show the Spearman's rho between the modules and each mOTU (*F. prausnitzii* 06108 on the x-axis and *F. prausnitzii* 06110 on the y-axis). Significant correlations (FDR-corrected q < 0.05) were kept and binned into KEGG modules. Red point denotes that the rho difference between the two mOTUs is larger 0.5.

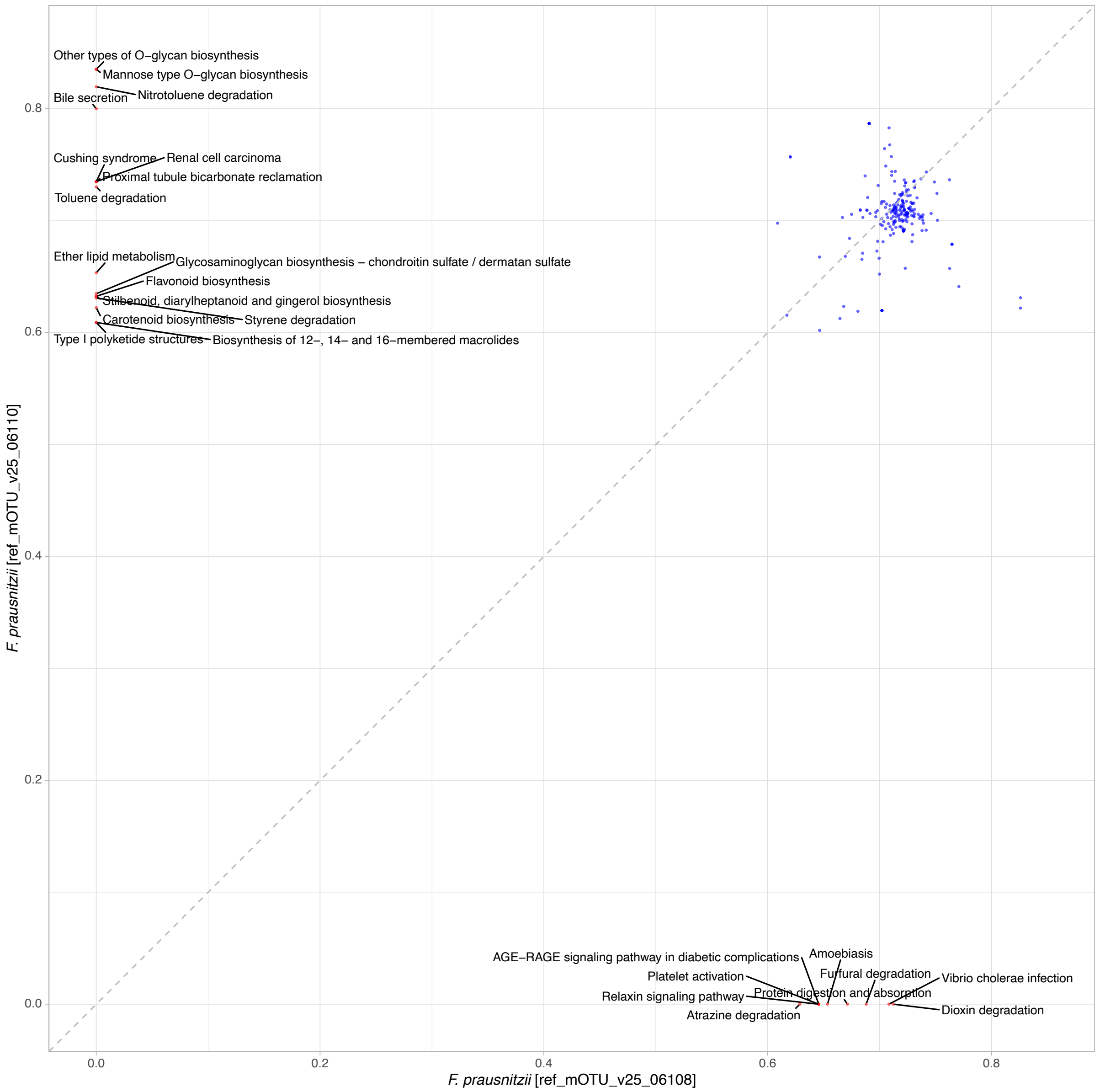


**Extended figure 8. The KEGG pathways that are correlated with the two uncultured *F. prausnitzii* mOTUs.** The x and y axes show the Spearman's rho between the pathways and each mOTU (*F. prausnitzii* 06108 on the x-axis and *F. prausnitzii* 06110 on the y-axis). Significant correlations (FDR-corrected q < 0.05) were kept and binned into KEGG pathways. Red point denotes that the rho difference between the two mOTUs is larger 0.5.

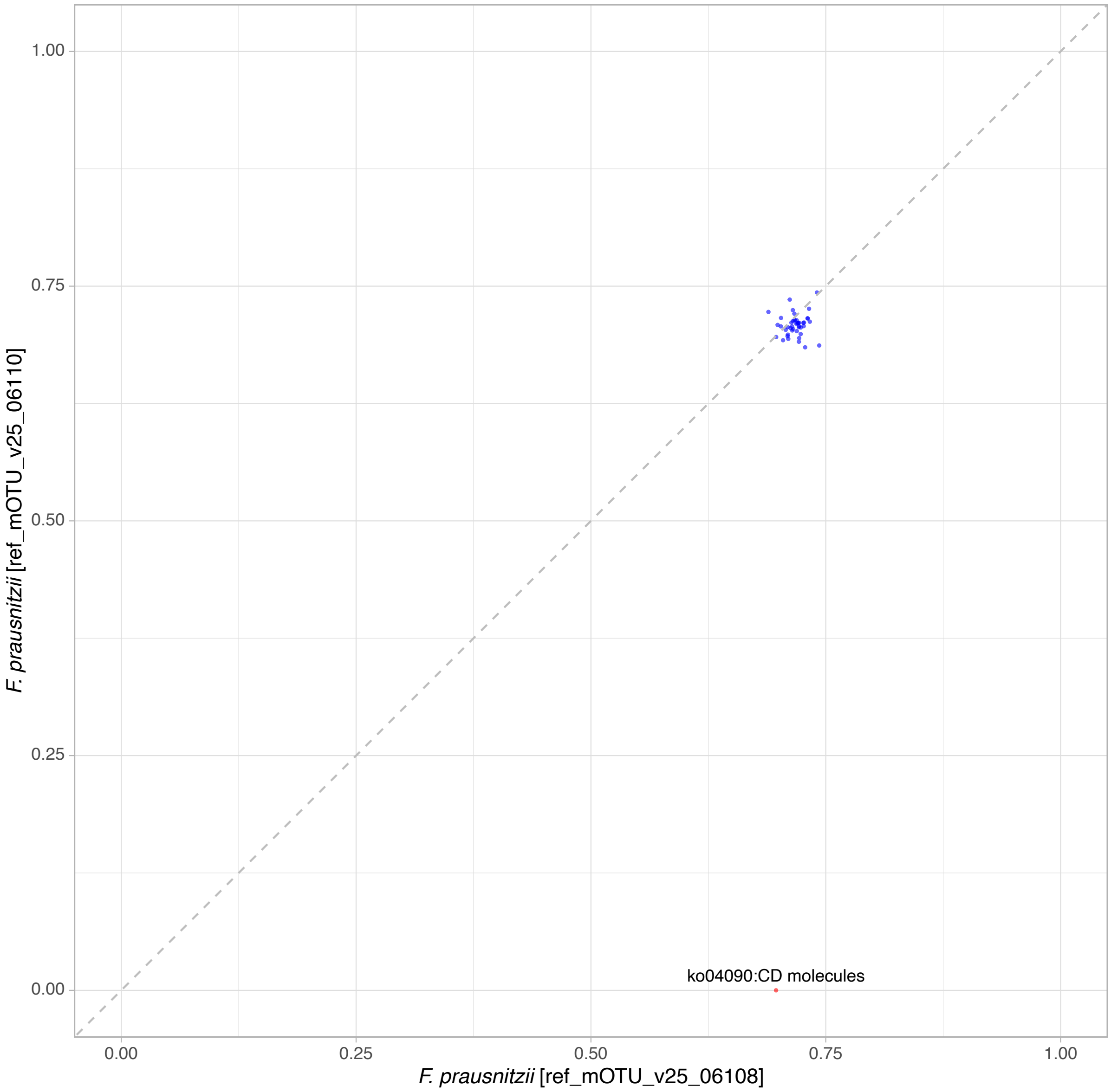


**Extended figure 9. The BRITEs that are correlated with the two uncultured *F. prausnitzii* mOTUs.** The x and y axes show the Spearman's rho between the BRITEs and each mOTU (*F. prausnitzii* 06108 on the x-axis and *F. prausnitzii* 06110 on the y-axis). Significant correlations (FDR-corrected q < 0.05) were kept and binned into BRITEs. Red point denotes that the rho difference between the two mOTUs is larger 0.5.
